# Class I and II aminoacyl-tRNA synthetase tRNA groove discrimination created the first synthetase•tRNA cognate pairs and was therefore essential to the origin of genetic coding

**DOI:** 10.1101/593269

**Authors:** Charles W. Carter, Peter R. Wills

## Abstract

The genetic code likely arose when a bidirectional gene began to produce ancestral aminoacyl-tRNA synthetases (aaRS) capable of distinguishing between two distinct sets of amino acids. The synthetase Class division therefore necessarily implies a mechanism by which the two ancestral synthetases could also discriminate between two different kinds of tRNA substrates. We used regression methods to uncover the possible patterns of base sequences capable of such discrimination and find that they appear to be related to thermodynamic differences in the relative stabilities of a hairpin necessary for recognition of tRNA substrates by Class I aaRS. The thermodynamic differences appear to be exploited by secondary structural differences between models for the ancestral aaRS called synthetase Urzymes and reinforced by packing of aromatic amino acid side chains against the nonpolar face of the ribose of A76 if and only if the tRNA CCA sequence forms a hairpin. The patterns of bases 1, 2 and 73 and stabilization of the hairpin by structural complementarity with Class I, but not Class II aaRS Urzymes appears to be necessary and sufficient to have enabled the generation of the first two aaRS•tRNA cognate pairs, and the launch of a rudimentary binary genetic coding related recognizably to contemporary cognate pairs. As a consequence, it seems likely that non-random aminoacylation of tRNAs preceded the advent of the tRNA anticodon stem-loop. Consistent with this suggestion, coding rules in the acceptor-stem bases also reveal a palimpsest of the codon•anticodon interaction, as previously proposed.

## Introduction: the aminacyl-tRNA synthetase duality; whence cognate pairs?

Carter and Wolfenden showed in 2015 that tRNA is nature’s repository for information about how amino acids drive protein folding (10, 42, 11). Embedding that information into tRNA required quasi-specific recognition of opposite grooves of ancestral tRNA minihelices by ancestral Class I and II aminoacyl-tRNA synthetases. In another recent publication (4), we noted that prior to the development of genetic coding coded sequence space was necessarily formless and void, until a bidirectional tRNA synthetase Protozyme gene (33) colonized two disjoint regions, coding for two distinct peptides with amino acid activating activity that each could use to select amino acids from its own Class, thereby implementing the reflexive feedback necessary to evolve and refine coding precision.

Converting the synthetase duality into the first two cognate pairs however, required a parallel duality in ancestral tRNA substrates, which has been quite difficult to discover. What differentiated the earliest Class I and Class II tRNA substrates? The tRNA acceptor stem bases that determine which Class of aaRS will acylate its terminal adenosine do not furnish any obvious clues as to how this determination might initially have been made. The frequencies of the four bases in positions 1, 2, 72, and 73 are nearly the same for Class I and II tRNA substrates (Fig. 1), making it difficult to discern patterns necessary to form the first aaRS•tRNA cognate pairs.

**Figure 1.**
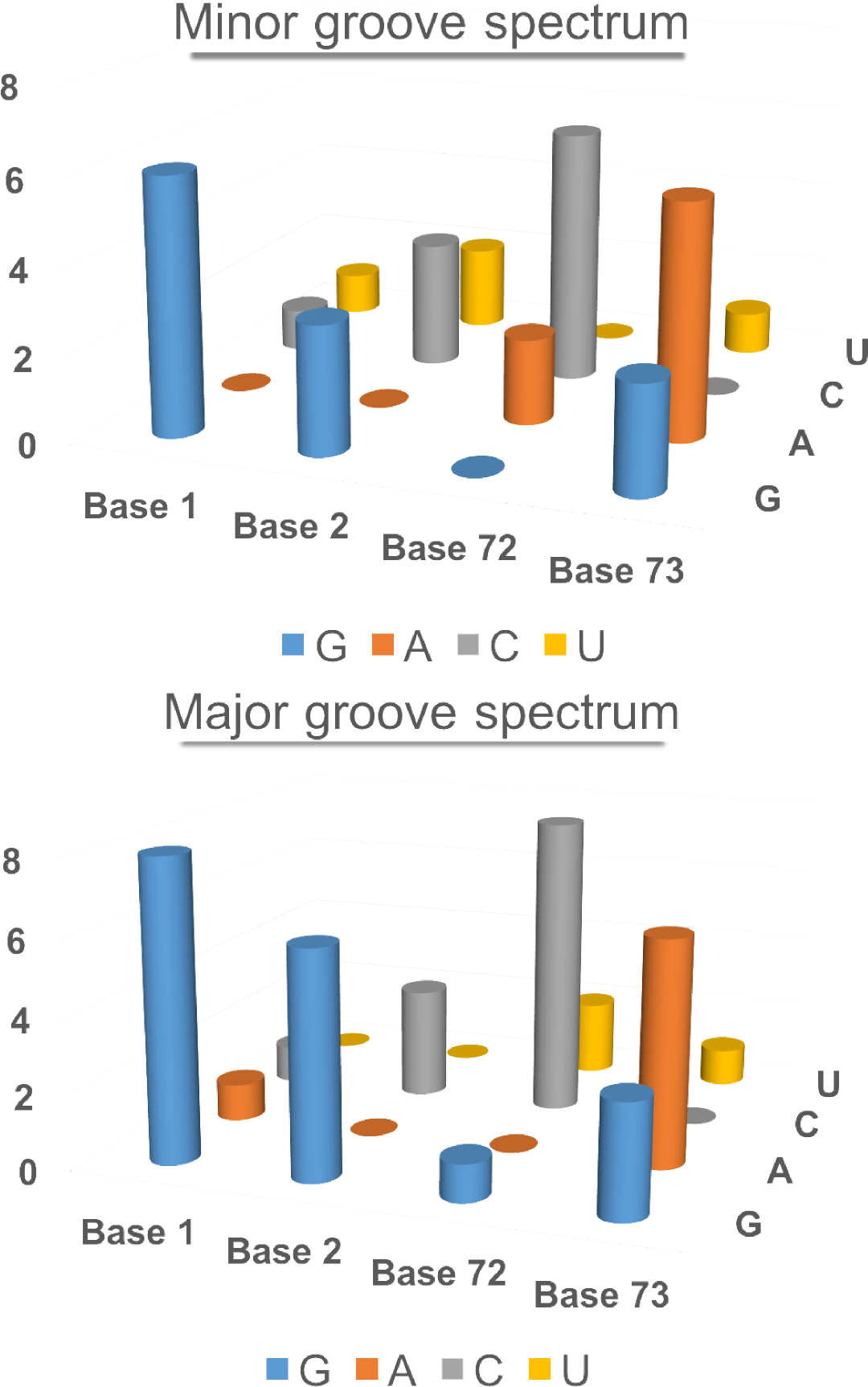
Base composition spectra of the topmost four bases of the acceptor stems of tRNAs recognized by their minor and major grooves. The two histograms almost superimpose, and have an R^2^ value of 0.79.

To address this question we used statistical regression methods, similar to those used to map the amino acid phase transfer free energies to the acceptor stem and anticodon (10, 42), to show that this primordial differentiation is coded in the three topmost tRNA acceptor stem bases (3). As new amino acids entered the code, they overwrote that ancestral tRNA duality, turning the acceptor-stem bases into a Palimpsest, derived from the Greek verb “psen”, “to scrape out”.

## The Operational RNA Code

Experimental acylation of tRNA minihelices by full-length aaRSs led Schimmel, Giegé, Moras, and Yokohama (39) to renew the earlier speculation (23) that aaRS catalytic domains and minihelices originally mediated protein synthesis according to a code residing entirely within the acceptor stem, which they called an “operational RNA code”. AARS Urzymes (36, 35, 31, 30, 7) are substantially smaller than catalytic domains, and cannot recognize tRNA anticodons, but nevertheless acylate intact tRNAs (30), validating their proposal.

Crystal structures for aaRS•tRNA complexes (37, 38, 25, 16, 45, 40, 44, 29) show that Class I synthetases approach from the minor groove, requiring a hairpin to insert the A76 ribose into the active site (Fig. 2). The acceptor stem is undistorted in tRNA complexes of Class II synthetases. Synthetases for aromatic amino acids are exceptional in that tRNA^Phe^ (25) does form the hairpin, whereas tRNA^Tyr^ (15) and tRNA^Trp^ (40, 44) do not.

**Figure 2.**
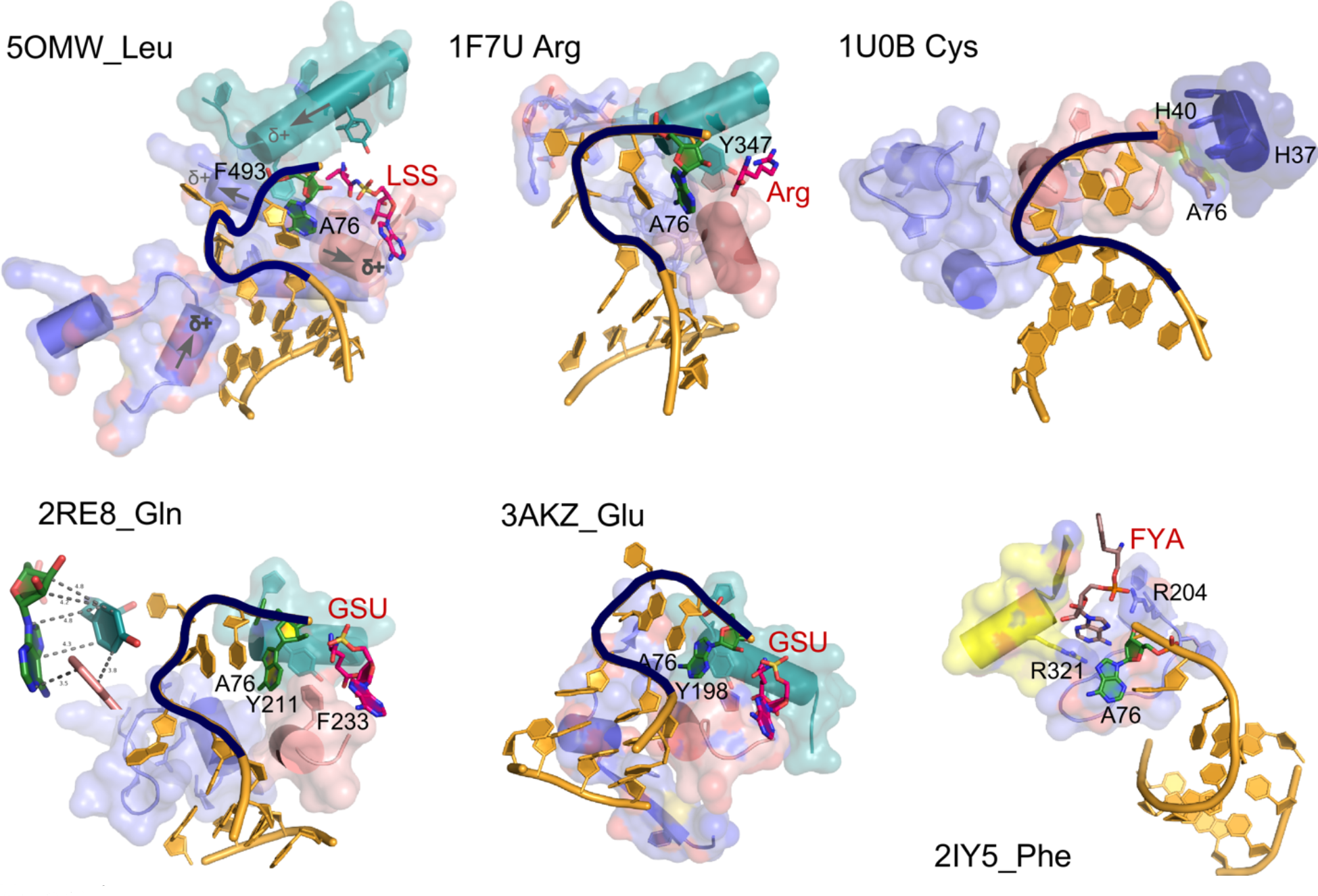
AARS•tRNA complexes interacting from the minor groove side. PDB IDs and amino acids are as indicated. CCA hairpin configurations are highlighted in black. Active-site ligands are abbreviated as follows: LSS, 5’-O-(L-leucylsulfamoyl)adenosine; Arg, arginine; GSU, O5’-(L-glutamyl-sulfamoyl)-adenosine; FYA, adenosine-5’-[phenylalaninol-phosphate]. Class I secondary structures depicted by color include a homologous section of connecting peptide 1 (CP1), slate; the “specificity-determining helix” teal; and the base of the helix from the second crossover of the Rossmann fold, containing the signature GxDQ, salmon), and in the case of PheRS the Motif-2 loop (slate) and Motif-3 (yellow). Electrostatic influences of the dipole moments of various helices are indicated on the helices of 5OMW_Leu. Side chains that interact with the tRNA 3’-terminal adenosine (A76) are indicated by number. Note that in the 1U0B (CysRS) complex the 3’-terminal adenosine occupies an unproductive position close to the HIGH catalytic signature, suggesting that in the absence of aminoacyl-5’-adenylate, the adenine ring finds a site similar to that normally occupied by the adenine ring of the adenylate, underscoring its potential binding affinity for the heterocycle. Inset in 2ER8_Gln shows the interaction between A76 and a conserved aromatic side chain at the N-terminus of the specificity-determining helix (adapted from (3), with permission).

## Multiple Regression Analysis

We assembled a design matrix with one row for each amino acid and columns for the groove recognized—the dependent variable—and two columns for each base, containing binary bits, (1 or –1) as predictors—the independent variables (Fig. 3). Multiple Regression uses least squares to solve the simultaneous equations implicit in assigning coefficients, β_i_, that best estimate the observed Groove in column 2, either –1 for the major or 1 for the minor groove. Each base is specified completely by two bits in the design matrix, providing the Predictors, P_i_. Because the entries of design matrix are digital, it can be viewed as a bar-code (Fig. 3C). The information in Fig. 3C is equivalent to that in Supplementary Table SI in Carter and Wills (3). The RHS of the regression equation combines an expression for the calculated value of the dependent variable with an error term, ε, representing the quantity to be minimized.

**Figure 3.**
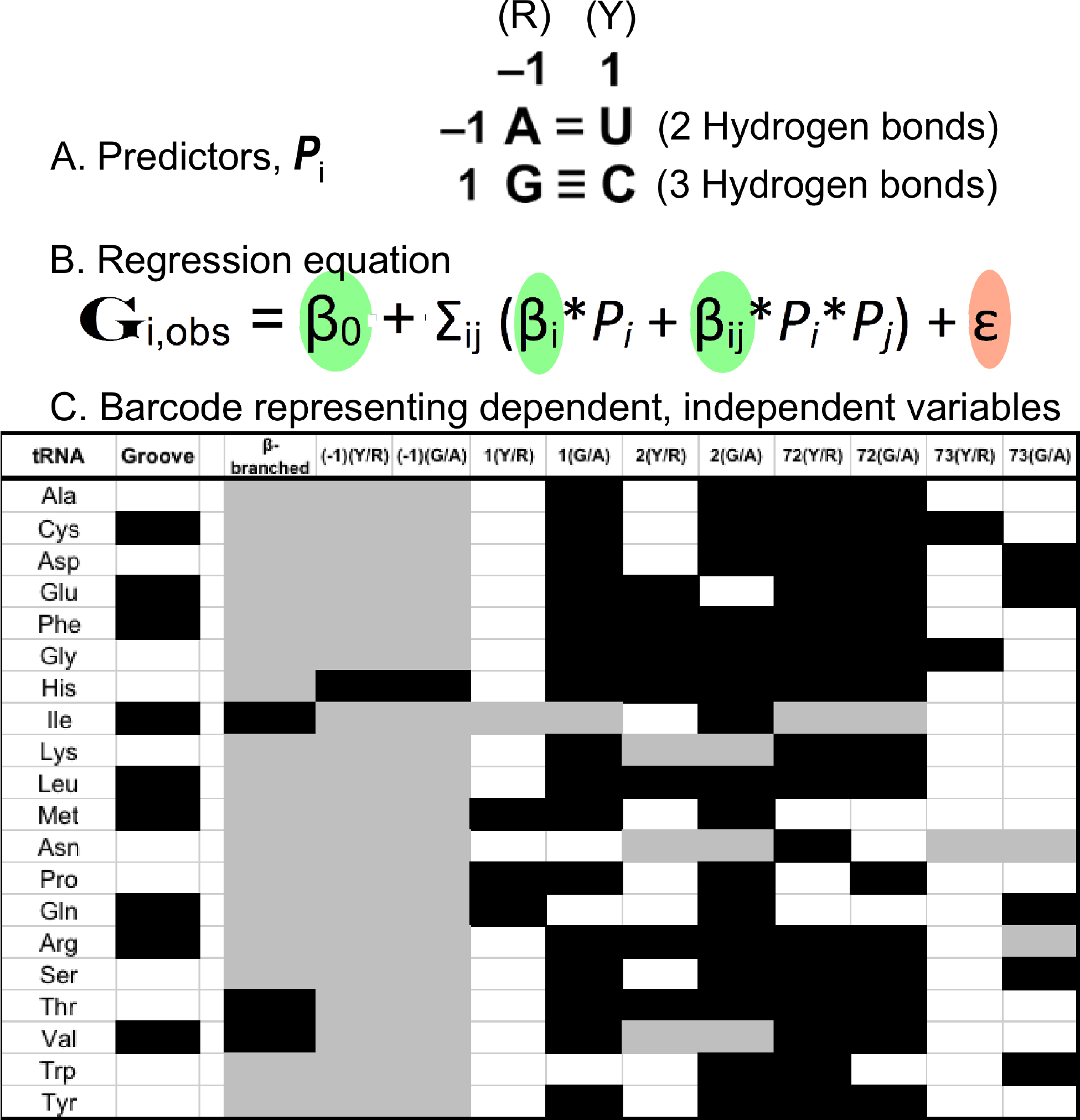
Regression modeling of tRNA groove recognition. A. Predictors are formulated by assigning two binary descriptors, {1, −1} to each base in the acceptor stem. A zero indicates that a predictor is not involved in groove recognition. B. The set linear vector equations relating the observed Groove, **G**_i_, to a linear combination computing the calculated groove value, plus an error, ε, which is the quantity minimized by least squares. C. Barcode representation of dependent, independent variables emphasizes the computational aspect of the acceptor-stem coding. Black, white, and grey denote values of {1}, {−1} and (13) respectively.

The corresponding design matrix based solely on ID elements (10) was much more sparsely populated than that used here. We repeatedly failed to find useful regression models for groove recognition using only the previously identified identity elements (24). Yet, the tRNA duality emerges directly from the more densely populated design matrix used here, (Fig. 3C). We were, in fact, led to include descriptors for the top two base pairs of all tRNA isoacceptors into the design matrix by analyzing outliers that couldn’t be fitted to regression models based on the sparser matrix. We also chose to analyze only bacterial tRNA sequences, to avoid corruption of any signal by the slight variations in tRNA identity in archaea and eukarya. The full model for all 20 isoacceptors has 11 coefficients. Remarkably, both least squares coefficients and the system of equations are entirely digital, so that the model predicts the correct groove identically for all 20 canonical amino acids (Table 1). Moreover, the fact that all sets of coefficients have integral or half-integral values reinforces the conclusion that this is a binary, digital, molecular code.

**Table 1.** Computation of G_calc_. Each cell underneath an amino acid shows the product of the estimate in the second column with the appropriate element of the design matrix in Table S1; totals are accumulated in the bottom row.

Class I
| Term | Estimate | Cys | Glu | Ile | Leu | Met | Gln | Arg | Val | Trp | Tyr |
| --- | --- | --- | --- | --- | --- | --- | --- | --- | --- | --- | --- |
| Intercept | 0.5 | -0.500 | -0.500 | -0.500 | -0.500 | -0.500 | -0.500 | -0.500 | -0.500 | -0.500 | -0.500 |
| β-branched | -2.5 | 0 | 0 | -2.500 | 0 | 0 | 0 | 0 | -2.500 | 0 | 0 |
| $(-1)(Y/R)$ | 2 | 0 | 0 | 0 | 0 | 0 | 0 | 0 | 0 | 0 | 0 |
| $1(Y/R)$ | 0.5 | -0.500 | -0.500 | 0 | -0.500 | 0.500 | 0.500 | -0.500 | -0.500 | -0.500 | -0.500 |
| $2(Y/R)$ | 1 | -1.000 | 1.000 | -1.000 | 1.000 | -1.000 | -1.000 | 1.000 | 0 | -1.000 | -1.000 |
| $2(G/A)$ | 1 | 1.000 | -1.000 | 1.000 | 1.000 | 1.000 | 1.000 | 1.000 | 0 | 1.000 | 1.000 |
| $72(G/A)$ | -0.5 | -0.500 | -0.500 | 0.000 | -0.500 | 0.500 | 0.500 | -0.500 | -0.500 | 0.500 | -0.500 |
| β-branched*$2Y/R$ | -4 | 0 | 0 | 4.000 | 0 | 0 | 0 | 0 | 0 | 0 | 0 |
| $2(Y/R)*73(Y/R)$ | -1 | 1.000 | 1.000 | -1.000 | 1.000 | -1.000 | -1.000 | 1 | 0 | -1.000 | -1.000 |
| $1(Y/R)*72(G/A)$ | -0.5 | 0.500 | 0.500 | 0 | 0.500 | 0.500 | 0.500 | 0.500 | 0.500 | -0.500 | 0.500 |
| $2(Y/R)*2(G/A)$ | -1 | 1.000 | 1.000 | 1.000 | -1.000 | 1.000 | 1.000 | -1.000 | 0 | 1.000 | 1.000 |
| β-branched*$72GA$ | 4.5 | 0 | 0 | 0 | 0 | 0 | 0 | 0 | 4.5 | 0 | 0 |
| Total |  | 1.000 | 1.000 | 1.000 | 1.000 | 1.000 | 1.000 | 1.000 | 1.000 | -1.000 | -1.000 |

Class II
| Term | Estimate | Ala | Asp | Phe | Gly | His | Lys | Asn | Pro | Ser | Thr |
| --- | --- | --- | --- | --- | --- | --- | --- | --- | --- | --- | --- |
| Intercept | -0.5 | -0.500 | -0.500 | -0.500 | -0.500 | -0.500 | -0.500 | -0.500 | -0.500 | -0.500 | -0.500 |
| β-branched | -2.5 | 0 | 0 | 0 | 0 | 0 | 0 | 0 | 0 | 0 | -2.500 |
| $(-1)(Y/R)$ | 2 | 0 | 0 | 0 | 0 | -2.000 | 0 | 0 | 0 | 0 | 0 |
| $1(Y/R)$ | 0.5 | -0.500 | -0.500 | -0.500 | -0.500 | -0.500 | -0.500 | -0.500 | 0.500 | -0.500 | -0.500 |
| $2(Y/R)$ | 1 | -1.000 | -1.000 | 1.000 | 1.000 | 1.000 | 0 | 0 | -1.000 | -1.000 | 1.000 |
| $2(G/A)$ | 1 | 1.000 | 1.000 | 1.000 | 1.000 | 1.000 | 0 | 0 | 1.000 | 1.000 | 1.000 |
| $72(G/A)$ | -0.5 | -0.500 | -0.500 | -0.500 | -0.500 | -0.500 | -0.500 | 0.500 | -0.500 | -0.500 | -0.500 |
| β-branched*$2Y/R$ | -4 | 0 | 0 | 0 | 0 | 0 | 0 | 0 | 0 | 0 | -4.000 |
| $2(Y/R)*73(Y/R)$ | -1 | -1.000 | -1.000 | 1.000 | -1.000 | 1.000 | 0 | 0 | -1.000 | -1.000 | 1.000 |
| $1(Y/R)*72(G/A)$ | -0.5 | 0.500 | 0.500 | 0.500 | 0.500 | 0.500 | 0.500 | -0.500 | -0.500 | 0.500 | 0.500 |
| $2(Y/R)*2(G/A)$ | -1 | 1.000 | 1.000 | -1.000 | -1.000 | -1.000 | 0 | 0 | 1.000 | 1.000 | -1.000 |
| β-branched*$72GA$ | 4.5 | 0 | 0 | 0 | 0 | 0 | 0 | 0 | 0 | 0 | 4.5 |
| Total |  | -1.000 | -1.000 | 1.000 | -1.000 | -1.000 | -1.000 | -1.000 | -1.000 | -1.000 | -1.000 |

## Hierarchical Rules: A Major Subset of Amino Acids Obeys a Very Simple Model

The coefficients imply rules determining groove recognition by aaRS. These are summarized in Table 2. The twenty amino acids can be partitioned into a set of 16 that obey a subset of the rules in Table 2 with a complement of 4 that require additional coefficients. The four outliers are histidine, for which the dominant discriminating feature is a G in the 5’ position, –1, plus the three amino acids with β-branched side chains. Thus, the set including Class I Leu, Met, Cys, Arg, Glu, Gln, Tyr, and Trp all obey the same subset of the rules in Table 3 with only seven coefficients plus the intercept as do Class II Ala, Asn, Asp, Gly, Lys, Ser, Pro, and Phe. That model therefore has 8 degrees of freedom, the same number as the model in Table 3. Moreover, the coefficients are identical for the two models, and the subset model gives R^2^ = 1.0.

**Table 2.** AARS groove recognition rules implied by the four topmost acceptor-stem bases.

| Term | Estimate |  | Implications |
| --- | --- | --- | --- |
| Intercept | -0.5 |  |  |
| β-branched | -2.5 | Coeff > 0; | β-branched=> minor groove<br>Unbranched=> major groove |
| -1,Y/R | 2 | Coeff > 0; | Base -1 purine => major groove<br>Base -1 pyrimidine => minor groove |
| 1,Y/R | 0.5 | Coeff > 0; | Base 1 pyrimidine => minor groove<br>Base 1 purine => major groove |
| 2,Y/R | 1 | Coeff > 0; | Base 2 pyrimidine => minor groove<br>Base 2 purine => major groove |
| 2,≡/= | 1 | Coeff > 0; | Base 2 G,C => minor groove<br>Base 2 A U => major groove |
| 72,≡/= | -0.5 | Coeff < 0; | Base 72 G/C => major groove<br>Base 72 A/U => minor groove |
| β-branched*2,Y/R | -4 | Coeff < 0; | Branched sc with Base 2 = C,U => major groove<br>Branched sc with Base 2 = G,A => minor groove |
| 1,Y/R*72,≡/= | -0.5 | Coeff < 0; | C=G => major groove;<br>G=C => minor groove<br>A=U => major groove<br>U=A => minor groove |
| 2,Y/R*73,Y/R | -1 | Coeff < 0; | Base 2 = C,U and Base 73=C,U => major groove<br>Base 2 = C,U Base 73 = G,A => minor groove<br>Base 2 = G,A Base 73 =C,U => minor groove<br>Base 2 = G,A Base 73 =G,A => major groove |
| 2,Y/R*2,≡/= | -1 | Coeff < 0; | **C => major groove**<br>**U => minor groove**<br>**G => minor groove**<br>**A => major groove** |
| β-branched*72,≡/= | 4.5 | Coeff > 0; | Branched sc with Base 72 G, C => minor Groove<br>Branched sc with Base 72 A, U => major Groove |

**Table 3.** Groove computation with 3 coefficients for a reduced amino acid set. This table is identical in intent to Table S2. Here, however, the bases themselves are also tabulated, together with the products of their coefficients and the corresponding element of the design matrix. Inspection of Figure 1 will confirm the entries in columns 3-7. The intercept for this model is 1, the coefficients are: −1(Y/R) = 2, 1(Y/R) = 1, and 2(Y/R)*73(Y/R) = −1, leading to the respective contributions in columns 9-11 to the total in column 12.

| Class I |  |  | Base | Base | Base | Base | Constant | -1 | 1Y/R | 2Y/R*73Y/R | Total |
| --- | --- | --- | --- | --- | --- | --- | --- | --- | --- | --- | --- |
| IA | Arg |  | G | C | C | G | 1 | 0 | -1 | 1 | 1 |
| IA | Cys |  | G | G | C | U | 1 | 0 | -1 | 1 | 1 |
| IA | Leu |  | G | U | C | G | 1 | 0 | -1 | 1 | 1 |
| IA | Met |  | U | G | A | G | 1 | 0 | 1 | -1 | 1 |
| IB | Glu |  | G | C | C | A | 1 | 0 | -1 | 1 | 1 |
| IB | Gln |  | C | G | A | A | 1 | 0 | 1 | -1 | 1 |
| IC | Tyr |  | G | G | C | A | 1 | 0 | -1 | -1 | -1 |
| IC | Trp |  | A | G | U | G | 1 | 0 | -1 | -1 | -1 |
| Class II |  |  |  |  |  |  |  |  |  |  |  |
| IIA | Ala |  | G | G | C | A | 1 | 0 | -1 | -1 | -1 |
| IIA | Gly |  | G | G | C | G | 1 | 0 | -1 | -1 | -1 |
| IIA | His | G | G | C | C | A | 1 | -2 | -1 | 1 | -1 |
| IIA | Ser |  | G | C | C | U | 1 | 0 | -1 | -1 | -1 |
| IIB | Asp |  | G | G | C | G | 1 | 0 | -1 | -1 | -1 |
| IIC | Phe |  | G | C | C | A | 1 | 0 | -1 | 1 | 1 |

An even simpler model for the major partition has only two coefficients: 1(Y/R) and 2(Y/R)*73(Y/R). The two predictors fit all but Lys, Asn, and Pro from within that set of 16 perfectly (R^2^ = 1.0; logWorth ~ 80) with digital coefficients, {1, 1, −1}. This model uses only three degrees of freedom to fit 13 tRNA isoacceptors, and thus has 10 degrees of freedom. Curiously, the three amino acids with aromatic side chains remain within the set predicted by this very simple model. The thirteen qualifying tRNA isoacceptors thus contain representatives from each of the three recognized subclasses in each Class (Fig. 4). Computation of groove recognition is given for this subset in Table 2. The intercept of this model is +1.0, which implies that minor groove recognition is the default for the subset.

**Figure 4.**
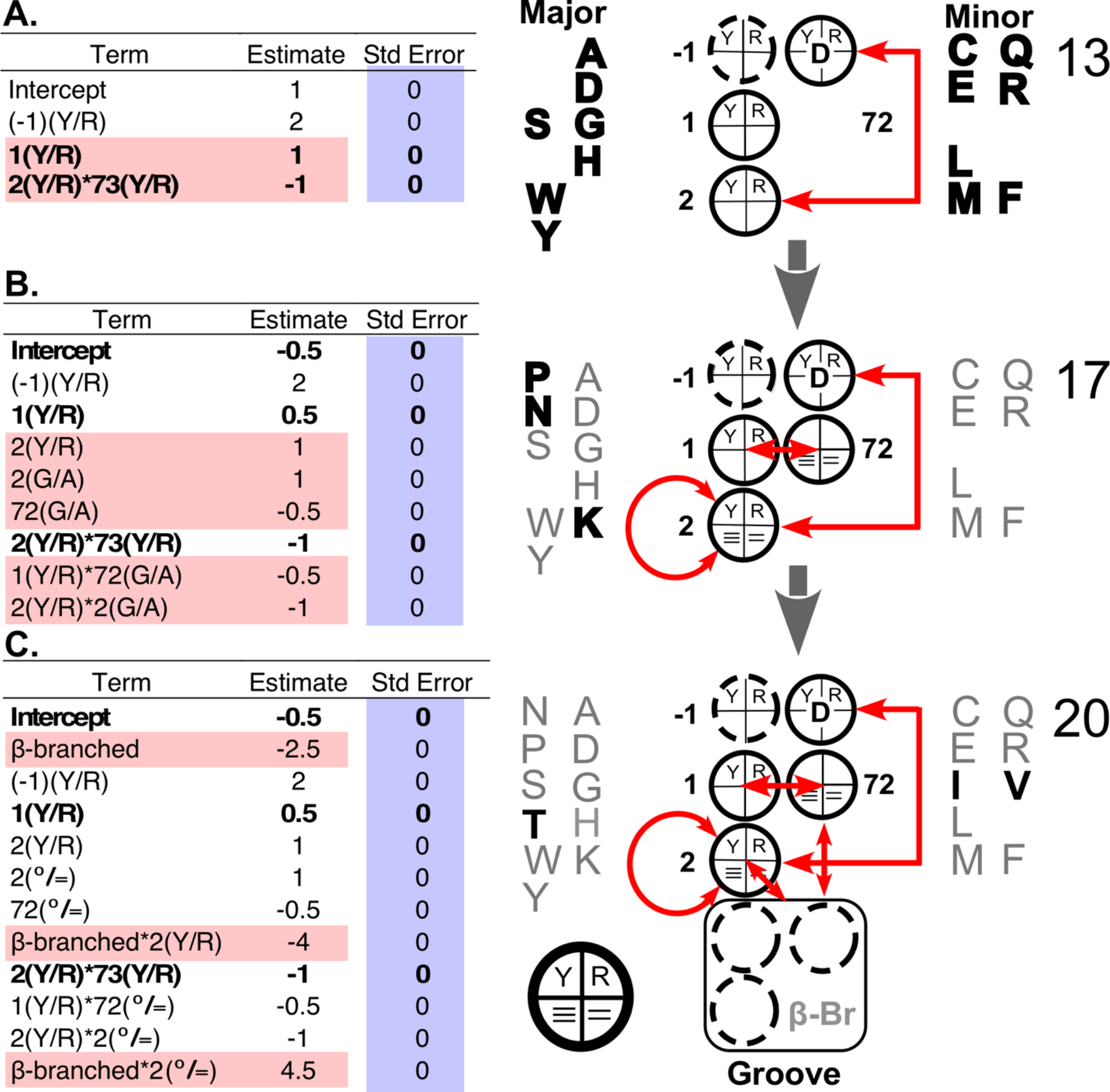
Hierarchy of models relating groove recognition to tRNA bases 1, 2, 72, 73. A. Minimal model for 13 isoacceptors. B. Intermediate model for 17 isoacceptors. C. Full model for 20 isoacceptors. (A), (B), and (C) each have three separate parts. On the left is a table showing model coefficients. The designator n(Y/R) refers to a binary variable whose value is {1} for a pyrimidine and {−1} for a purine at the base given by the number. The designator n(≡/=) refers to a binary variable whose value is {1} for either (G,C) and {−1} for (A,U). New coefficients in the hierarchy are outlined in red. Standard errors are highlighted in blue. The standard error of 0 means that the t-test probability of predicting the relevant grooves as well using random coefficients is <~10^−60^. The second column of the figure consists of schematic diagrams for the models, the key at the bottom right of C shows how the two bits for each base are used to form rules. Red arrows denote two-way interactions.

A surprising hierarchy appears in the groove recognition rules if we eliminate rows of design matrix (Fig. 3C) corresponding to the outliers. Two coefficients suffice to predict 13 rows of isoacceptors. Eight coefficients predict an intermediate subset of 17 rows, excepting only branched side chains. None of those coefficients change as the β-branched amino acids are added. Rather, different values of the same coefficients are necessary to predict acceptor stem sequences for tRNA isoacceptors for β-branched amino acids.

## Groove Recognition Rules Imply Differential Stability of the helical CCA extension

The simple 2-coefficient model (Table 3) correctly predicts the groove recognized by 13 of the aaRS. Remarkably, it distinguishes only between pyrimidines and purines, suggesting that a thermodynamic basis for hairpin formation (Fig. 2) may underlie the original discrimination necessary to create synthetase•tRNA cognate pairs:

Rule (1): A base 1 pyrimidine implies the minor groove; a purine the major groove.
Rule (2): Both purines or both pyrimidines at Bases 2 and 73 imply the major groove; opposite base types imply the minor groove.

Values of base-stacking free energies from the literature (22, 43, 1) and from Mfold (46) both suggest that terminal base stacking in Class I tRNA substrates destabilizes the 3’ CCA extension by ~1.2 kcal/mole, enabling, but not necessarily mandating, hairpin formation. That modest thermodynamic distinction of Class I and II aaRS tRNA substrates, implied by the recognition rules (1) and (2), suggests that although tRNA bases 1, 2, 72, and 73 define aaRS Class recognition, the discrimination necessary to form the earliest operational aaRS•tRNA cognate pairs must also have had a component from the structures of the ancestral synthetases themselves.

## AARS Structures *Reinforce* the Rules

Urzyme structures inferred from extant crystal structures suggest that ancestral secondary structural differences between the two synthetase classes do actually reinforce the acceptor-stem structures observed in productive co-crystals (3). The Class I Urzyme Rossmann-fold architecture creates a positively-charged pocket for the A76 5’ phosphate at the N-terminus of the “specificity-determining helix” (Fig. 5). A conserved aromatic sidechain at the second position of the same helix packs against the hydrophobic faces of the ribose and adenine moieties, complementing that electrostatic dipole. Class I Urzymes thus provide a specific pocket for A76 if and only if it assumes a hairpin.

**Figure 5.**
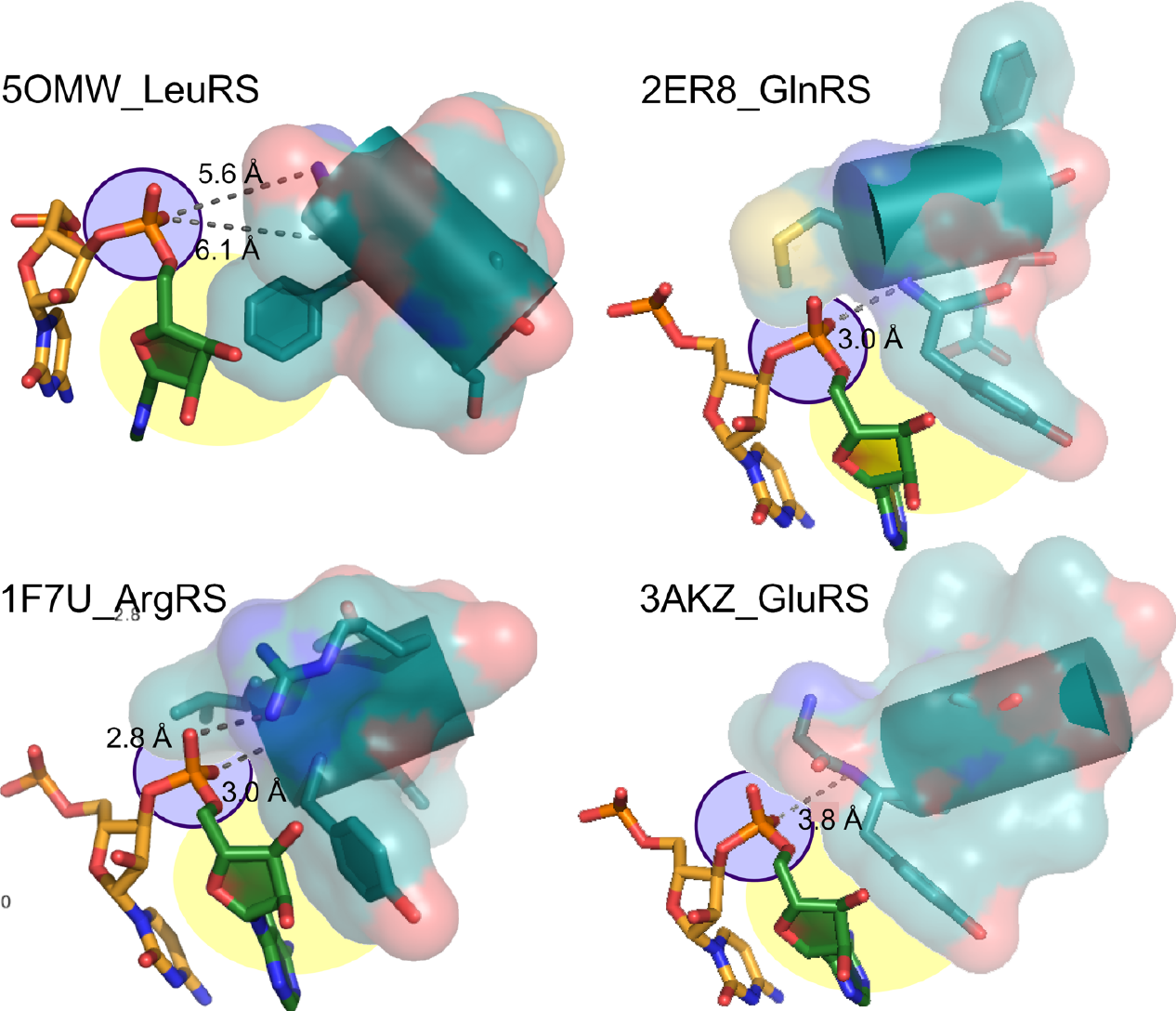
Interactions of Class I aaRS with the phosphate group of the 3’-terminal adenosine. The hairpin structure of Class I isoaccepting tRNA substrates orients this phosphate group so that it points toward the N-terminus of the specificity-determining helix (purple). Hydrogen bond distances shown suggest that, except in the case of 5OMW_LeuRS, these interactions may be strong. In 1F7U_ArgRS, the interaction is reinforced by a salt bridge between the phosphate group and R350. Specificity is enhanced by the stacking interaction between the nonpolar face of the A76 ribose and a conserved aromatic sidechain immediately following the four unpaired amide nitrogens at the N-terminus of the specificity-determining helix (yellow) (adapted from (3), with permission) (adapted from (3), with permission).

The Class II Motif 2 loop effectively protects the helical 3’ CCA extension by sitting on top of the first acceptor stem base pair (Fig. 6). Antiparallel β-structures have minimal dipole moments, and cannot perturb the RNA helix. Moreover, the β-structure is antiparallel to the 5’-3’ direction of the 3’ terminus, as predicted for such interactions (8). Notably, although these interactions are reinforced by conserved amino acid side chains, they arise fundamentally from complementary protein and RNA secondary structures whose determinants are independent of specific side chains, and therefore could have begun to function before genetic coding developed a full, precise alphabet.

**Figure 6.**
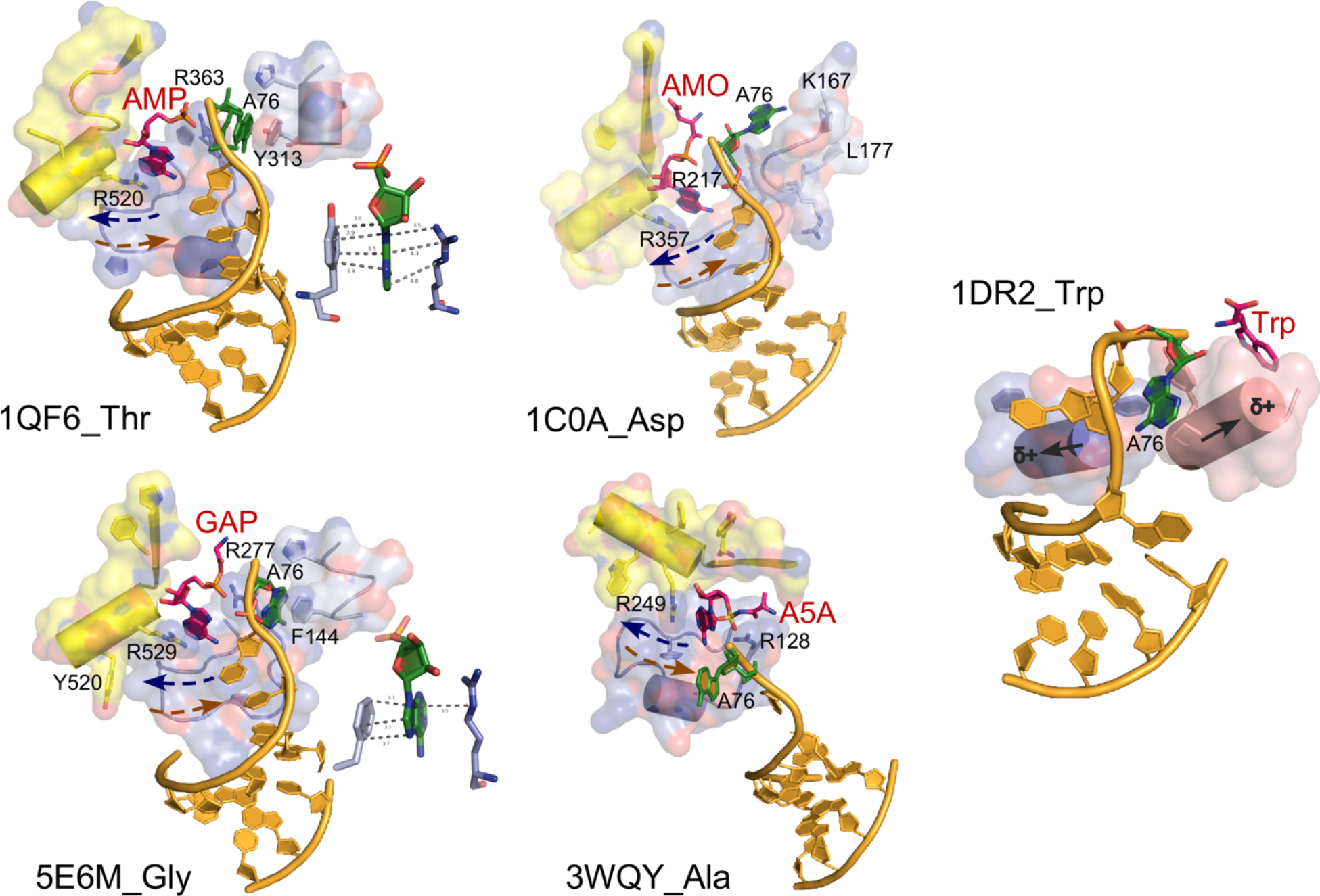
AARS•tRNA complexes interacting from the major groove side. PDB IDs and amino acids are as indicated. Active-site ligands are abbreviated as follows: AMP, adenosine 5’ monophosphate; GAP, glycyl-adenosine-5’-phosphate; AMO, aspartyl-adenosine-5’-monophosphate; Trp, L-tryptophan. Class II secondary structures depicted by color include the Motif-2 loop (slate), Motif-3 (yellow), and a loop that occurs prior to Motif-2, and in the case of TrpRS, the base of the helix from the second crossover of the Rossmann fold (salmon) and a segment of CP1 (slate). Side chains that interact with the tRNA 3’-terminal adenosine (A76) are indicated by number. Antiparallel helix directions of the Motif 2 loop and the 3’-CCA terminus are shown as dashed arrows on the four Class II complexes, 1QF6_Thr, 5EM6_Gly, 1C0A_Asp, and 3WQY_Ala. Chain directions indicated by the arrows in each case match that predicted by the Carter & Kraut model (8) for fitting antiparallel β-structure into the minor groove of A-form RNA. Helix dipoles indicated for 2DR2_Trp suggest that the reorientation of the tRNA^Trp^ acceptor stem has moved it to a position that minimizes electrostatic disruption of the RNA double helix. Insets in 1QF6_Thr and 5E6M_Gly show interactions between A76 and a conserved aromatic side chain N-terminal to the Motif 2 loop (adapted from (3), with permission).

The AlaRS complex is an outlier. The Motif II loop assumes approximately the same conformation as the other Class II aaRS, but is separated from the topmost base pairs by unwinding of the 3’ extension, to allow other synthetase domains not shown in Fig. 6 to recognize the G3-U70 base pair from both grooves (14). We have argued (3) that this modification of the synthetase•tRNA complex involves more recently acquired domains, and hence that it evolved more recently than the early expansion of the genetic code.

## Codon-Anticodon Pairing in the acceptor stem

Table 3 compiles the coding rules implied by the coefficients of the full model in Fig. 4C. The two bits defining base 2 are of special interest because they coordinately assign a specific groove unambiguously to each base. The two pyrimidines imply opposite grooves, as do those for the two purines. These implications coincide exactly with the coding specificity of the first two columns of the codon table and closely with the remaining columns.

Unexpectedly, that coincidence validates Di Giulio’s hypothesis that the acceptor stem retains a vestige of the codon-anticodon interaction (17–20). The anticodon in contemporary tRNAs is simply too far from the acceptor stem to permit such recognition by ancestral aaRS Urzymes. The operational RNA code in the acceptor stem almost certainly approximates a solution to that problem. Moreover, it would have allowed acylated minihelices to interact with proto-mRNAs before the advent of the anticodon loop.

## The operational RNA code

Discontinuity in the functional transfer of information via codon•anticodon pairing now appears more certain in view of this unexpected evidence for Di Giulio’s original proposal that the anticodon descended from sequences just prior to the 3’ CCA terminus of the acceptor stem. Such a discontinuity is also consistent with our discovery that the tRNA acceptor stem bases code for size, carboxylate vs non-carboxylate, and β-branched vs non-β-branched side chains (10, 11). Those coding distinctions now seem likely to have encoded much of the simplest and most ancestral coded proteins, including the earliest aaRS Protozymes and early ribosomal proteins, leaving the fine tuning of protein 3D structure to be resolved by the coding of polarity by the three anticodon bases.

The complete operational RNA code is a complex pattern of rules that differentiate 5 overlapping groups of isoacceptors, mapped here onto the acceptor stem bases (Fig. 7). Red arrows indicate two-way interactions, or energetic coupling between different binary bits associated with the same or different bases. The overlapping is very likely an important reason why these patterns are so elusive. Rules for groove and size are interdependent because the size model includes the groove. The others differ more markedly, and their origin extends further down the acceptor stem. For this reason, we previously referred to the difficult problem of sorting out the operational code as one of deciphering a palimpsest (3).

**Figure 7.**
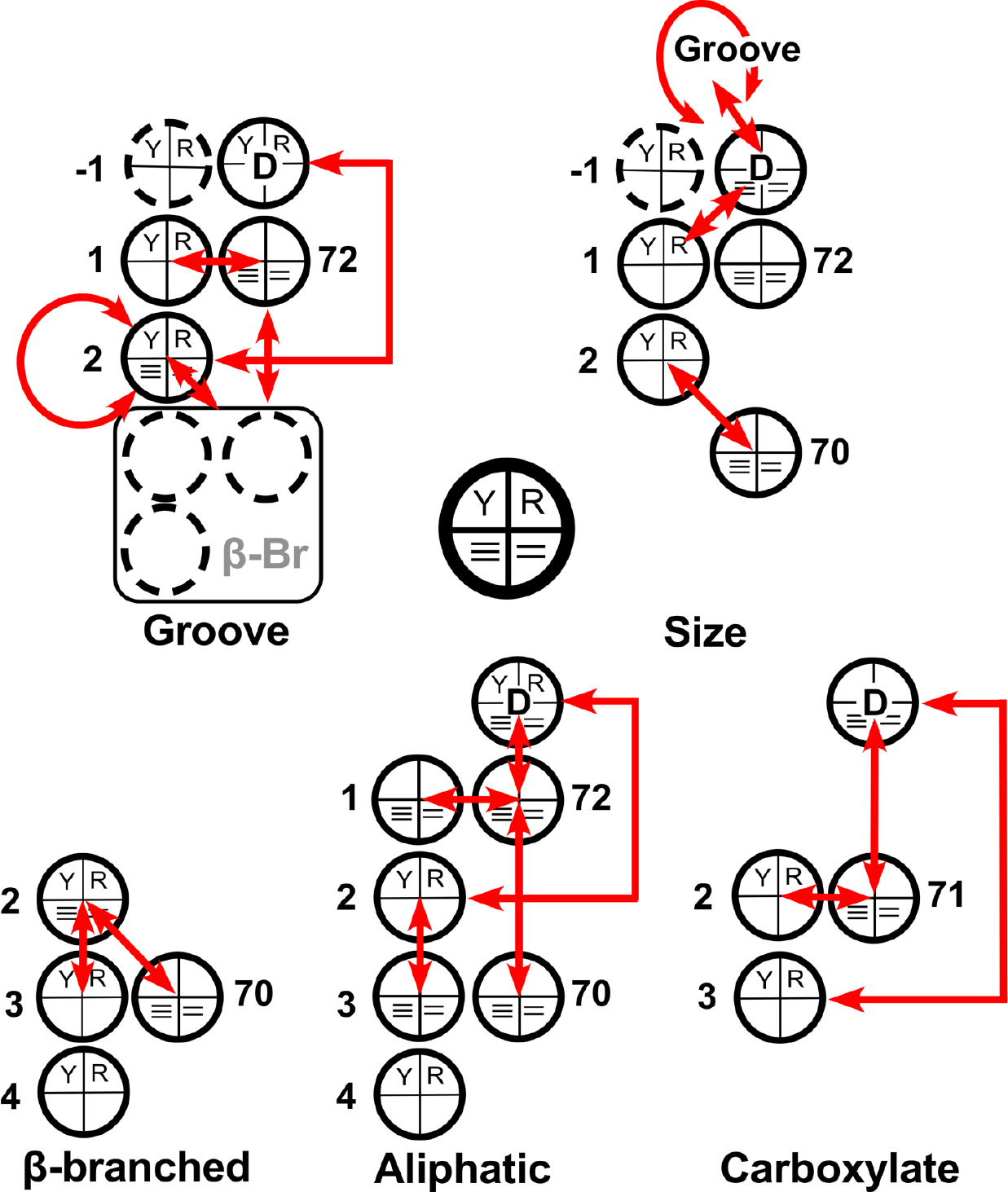
Elements of the “Operational RNA Code” (10, 11). Coding properties of the tRNA acceptor stem are represented schematically, to illustrate graphically relationships from each regression model in Tables 3 and 4. Bases that do not contribute to the respective regression model are omitted. Bases are represented by the schematic key, whose interpretation is given in Figure 3, which also include the regression coefficients (adapted from (3), with permission).

## Conclusions, future directions

The phylogenetic (34, 12, 6, 7, 2) and biochemical (36, 35, 31, 30, 5, 9, 7) evidence is quite strong that amino acid activation and tRNA aminoacylation began with the advent of ancestral synthetases with scarcely non-random specificities. A simple and consistent scenario (3) holds that the initial differentiation between amino acid types was that based on the sizes of their side chains, ancestral Class I and Class II protozymes activating amino acids with large and small side chains, respectively. For that conclusion to have been viable, a simultaneous and comparably simple basis must have existed for the original discrimination via synthetase-tRNA recognition, on which to form the first two synthetase•tRNA cognate pairs. It seems reasonable, therefore, that the differentiation of tRNA acceptor stems (or minihelices) occurred approximately simultaneously with the appearance of a bidirectional genetic quasispecies (exemplified by 33) encoding ancestral Class I and II Urzymes.

Groove recognition is dictated, surprisingly, by just the 4 topmost bases of the ancestral tRNA acceptor stem, which facilitate a hierarchical set of “rules” (Table 3). In this work (3) we have reviewed evidence that the original cognate pairs resulted from a combination of thermodynamic conformational preferences induced by patterns of acceptor stem purines and pyrimidines in the topmost four bases that enable hairpin formation by tRNA acceptor stems of ancestral Class I aaRS, and Class-specific synthetase secondary structures in both Classes that reinforced the resulting differential thermodynamic tendencies.

β-branched amino acids obey different Groove-recognition rules, that are essentially offset by changes in the constant terms that allow the remaining coefficients to function without change. The manner in which this is accomplished suggests that more remains to be deciphered concerning how these amino acids were recruited into the growing codon table. However, we have not been able to identify other correlations that might further illuminate this question.

Codon-anticodon pairing in the acceptor stem may have allowed coded translation to begin before tRNAs acquired anticodons, along lines long advocated by DiGiulio (17–20). Use of that pairing may also reflect a transitional mechanism for bidirectional coding via formation of triple helices (3).

Evolution of the universal genetic code appears to have recapitulated the rules for protein folding. The simplicity of the four choices of amino acid types enabled by the operational code (Figure 6) and the fact that some of the rules evident from the analysis are actually buried in contemporary acceptor-stem sequences suggest a temporal order to the appearance and evolution of secondary and tertiary structures. The initial two codewords for large and small amino acid side chains enabled the search for proto-mRNAs with binary patterns (27), which may already have begun to discriminate between extended β– and α–helical structures. That distinction would have been reinforced by adding selections for β-branched and aliphatic versus polar side chains, leading to a finer distinction between the two types of chain segments. Adding the anticodon, with its new precision with its newly created ability to specify side-chain polarity would then have enabled a much broader search for different tertiary folds (10, 11), stabilizing tertiary structure by tuning the differences between core and surface residues, and thus continuing to shape the evolution of protein structures.

There is an appealing continuity in this model for codon development, arising initially from aaRS groove discrimination in the acceptor stem. Characteristics of the operational code are consistent with recent suggestions that protein structures evolved coordinately with the alphabet size and that the evolution of the genetic code also traces the discovery of protein folding rules (4). Correlations between secondary and tertiary structures of ribosomal proteins and the estimated order of rRNA additions provoked a similar hypothesis from an entirely different direction (28). Those authors postulated staged protein evolution, earlier stages marked initially by the absence of secondary structures, followed by the presence of β-, then α-structures, and then only after a prolonged development of secondary structures, by the emergence of tertiary structures. Their hypothesis assumes co-evolution of protein folding with the genetic code, but makes no reference to how properties of the genetic code might have undergone comparable development. This work furnishes a necessary complement and rationalization for such observations.

An important application of this work is that joint synthetase/tRNA duality may constrain ancestral reconstruction sufficiently to recover simpler alphabets via conditioned ancestral sequence reconstructions (32, 26, 21), which now have significantly greater potential to identify confidently some or all of the reduced alphabets leading to the contemporary canonical twenty amino acids. We anticipate that recent structure-based curation of the aaRS sequence database (41) will provide additional confidence in such reconstructions.

## References

1 Brown R.F., Andrews C.T., and Elcock A.H. (2015) Stacking Free Energies of All DNA and RNA Nucleoside Pairs and Dinucleoside Monophosphates Computed Using Recently Revised AMBER Parameters and Compared with Experiment. J Chem Theory Comput 11, 2315–2328.

2 Carter C.W., Jr (2018) The Evolution of Genetic Coding. Scientia In press.

3 Carter C.W., Jr. and Wills P.R. (2018) Hierarchical groove discrimination by Class I and II aminoacyl-tRNA synthetases reveals a palimpsest of the operational RNA code in the tRNA acceptor-stem bases. Nucleic Acids Research 46, 9667–9683.

4 Carter C.W., Jr. and Wills P.R. (2018) Interdependence, Reflexivity, Fidelity, and Impedance Matching, and the Evolution of Genetic Coding. Molecular Biology and Evolution 35, 269–286.

5 Carter C.W., Jr. (2014) Urzymology: Experimental Access to a Key Transition in the Appearance of Enzymes. J Biol Chem 289, 30213–30220.

6 Carter C.W., Jr. (2016) An Alternative to the RNA World. Natural History 125, 28–33.

7 Carter C.W., Jr. (2017) Coding of Class I and II aminoacyl-tRNA synthetases. Advances in Experimental Medicine and Biology: Protein Reviews 18, 103–148.

8 Carter C.W., Jr. and Kraut J. (1974) A Proposed Model for Interaction of Polypeptides with RNA. Proceedings of the National Academy of Sciences, USA 71, 283–287.

9 Carter C.W., Jr., Li L., Weinreb V., Collier M., Gonzales-Rivera K., Jimenez-Rodriguez M., Erdogan O., and Chandrasekharan S.N. (2014) The Rodin-Ohno Hypothesis That Two Enzyme Superfamilies Descended from One Ancestral Gene: An Unlikely Scenario for the Origins of Translation That Will Not Be Dismissed. Biology Direct 9, 11.

10 Carter C.W., Jr. and Wolfenden R. (2015) tRNA Acceptor-Stem and Anticodon Bases Form Independent Codes Related to Protein Folding. Proc Nat Acad Sci USA 112 7489–7494.

11 Carter C.W., Jr.. and Wolfenden R. (2016) Acceptor-stem and anticodon bases embed amino acid chemistry into tRNA. RNA Biology 13, 145–151.

12 Chandrasekaran S.N., Yardimci G., Erdogan O., Roach J.M., and Carter C.W., Jr (2013) Statistical Evaluation of the Rodin-Ohno Hypothesis: Sense/Antisense Coding of Ancestral Class I and II Aminoacyl-tRNA Synthetases. Mol Biol Evol 30, 1588–1604.

13 Cheong H.-K., Kim N.-K.n., and Cheong C. (2015) RNA Structure: Tetraloops. Chichester: eLS. John Wley & Sons, Ltd.

14 Chong Y.E., Guo M., Yang X.-L., Kuhle B., Naganuma M., Sekine S.-i., Yokoyama S., and Schimmel P. (2018) Distinct ways of G:U recognition by conserved tRNA binding motifs. Proc Nat Acad Sci USA 115, 7527–7532.

15 Cusack S., Tukalo M., and Yaremchuk A. (2002) Class I tyrosyl-tRNA synthetase has a class II mode of cognate tRNA recognition. EMBO J 21, 3829–3840.

16 Delagoutte B., Moras D., and Cavarelli J. (2000) tRNA aminoacylation by arginyl-tRNA synthetase: induced conformations during substrates binding. EMBO J 19, 5599–5610.

17 Di Giulio M. (1989) Some Aspects of the Organization and Evolution of the Genetic Code. J Mol Evol 29, 191–201.

18 Di Giulio M. (1992) On the Origin of the Transfer RNA Molecule. J Theor Biol 159, 199–214.

19 Di Giulio M. (2004) The origin of the tRNA molecule:implications for the origin of protein synthesis. Journal of Theoretical Biology 226, 89–93.

20 Di Giulio M. (2009) A Comparison Among the Models Proposed to Explain the Origin of the tRNA Molecule: A Synthesis. J Mol Evol 69, 1–9.

21 Drummond A.J., Suchard M.A., Xie D., and Rambaut A. (2012) Bayesian phylogenetics with BEAUti and the BEAST 1.7. Mol Biol Evol 29, 1969–1973.

22 Freier S.M., Kierzek R.D., Jaeger O.A., Sugimoto N., Caruthers M.H., Neilson T., and Turner D.H. (1986) Improved free-energy parameters for predictions of RNA duplex stability. Proc Nati Acad Sci USA 83, 9373–9377.

23 Giegé R. (1972) Recherches sur la spécificité de reconnaissance des acides ribonucléiques de transfert par les aminoacyl-tRNA synthétases [Study on the specificity of recognition of transfer ribonucleic acids by aminoacyl-tRNA synthetases]. In: Biological Chemistry Strasbourg, France: Université Louis Pasteur.

24 Giegé R., Sissler M., and Florentz C. (1998) Universal rules and idiosyncratic features in tRNA identity. Nucleic Acids Res 26, 5017–5035.

25 Goldgur Y., Mosyak L., Reshetnikova L., Ankilova V., Lavrik O., Khodyreva S., and Safro M. (1997) The crystal structure of phenylalanyl-tRNA synthetase from thermus thermophilus complexed with cognate tRNAPhe. Structure 5, 59–68.

26 Hanson-Smith V., Kolaczkowski B., and Thornton J.W. (2010) Robustness of Ancestral Sequence Reconstruction to Phylogenetic Uncertainty. Mol Biol Evol 27, 1988–1999.

27 Kamtekar S., Schiffer J.M., Xiong H., Babik J.M., and Hecht M.H. (1993) Protein Design by Binary Patterning of Polar and Non-polar Amino Acids. Science 262, 1680–1685.

28 Kovacs N.A., Petrov A.S., Lanier K.A., and Williams L.D. Frozen in Time: The History of Proteins. Mol Biol Evol 34, 1252–1260.

29 Larson E.T., Kim J.E., Zuckera F., Kelley A., Mueller N., Napulib A.J., Verlinde C.L.M.J., Fan E., Buckner F.S., Van Voorhis W.C., Merritt E.A., and Hol W.G.J. (2010) Structure of Leishmania major Methionyl-tRNA Synthetase in Complex with Intermediate Products Methionyladenylate and Pyrophosphate. Biochimie In Press.

30 Li L., Francklyn C., and Carter C.W., Jr (2013) Aminoacylating Urzymes Challenge the RNA World Hypothesis. J Biol Chem 288, 26856–26863.

31 Li L., Weinreb V., Francklyn C., and Carter C.W., Jr (2011) Histidyl-tRNA Synthetase Urzymes: Class I and II Aminoacyl-tRNA Synthetase Urzymes have Comparable Catalytic Activities for Cognate Amino Acid Activation. J Biol Chem 286, 10387–10395.

32 Liberles D.A. (2007) Ancestral Sequence Reconstruction. Oxford: Oxford University Press.

33 Martinez L., Jimenez-Rodriguez M., Gonzalez-Rivera K., Williams T., Li L., Weinreb V., Chandrasekaran S.N., Collier M., Ambroggio X., Kuhlman B., Erdogan O., and Carter C.W.J. (2015) Functional Class I and II Amino Acid Activating Enzymes Can Be Coded by Opposite Strands of the Same Gene. J Biol Chem 290, 19710–19725.

34 O’Donoghue P.. and Luthey-Schulten Z. (2003) On the Evolution of Structure in Aminoacyl-tRNA Synthetases. Microbiol Mol Biol Rev 67, 550–573.

35 Pham Y., Kuhlman B., Butterfoss G.L., Hu H., Weinreb V., and Carter C.W., Jr (2010) Tryptophanyl-tRNA synthetase Urzyme: a model to recapitulate molecular evolution and investigate intramolecular complementation. J Biol Chem 285, 38590–38601.

36 Pham Y., Li L., Kim A., Erdogan O., Weinreb V., Butterfoss G., Kuhlman B., and Carter C.W., Jr (2007) A Minimal TrpRS Catalytic Domain Supports Sense/Antisense Ancestry of Class I and II Aminoacyl-tRNA Synthetases. Mol Cell 25, 851–862.

37 Rould M.A., Perona J.J., Söll D., and Steitz T., A. (1989) Structure of the *E. coli* Glutaminyl-tRNA Synthetase Complexed with tRNA^Gln^ and ATP at 2.8 Å Resolution. Science 246, 1135–1142.

38 Ruff M., Krishnaswamy S., Boeglin M., Poterszman A., Mitschler A., Podjarny A., Rees B., Thierry J.C., and Moras D. (1991) Class II Aminoacyl Transfer RNA Synthetases: Crystal Structure of Yeast Aspartyl-tRNA Synthetase Complexed with tRNA^Asp^. Science 252, 1682–1689.

39 Schimmel P., Giegé R., Moras D., and Yokoyama S. (1993) An operational RNA code for amino acids and possible relationship to genetic code. Proc Nat Acad Sci USA 90, 8763–8768.

40 Shen N., Guo L., Yang B., Jin Y., and Ding J. (2006) Structure of human tryptophanyl-tRNA synthetase in complex with tRNA(Trp) reveals the molecular basis of tRNA recognition and specificity. Nucleic Acids Res 34, 3246–3258.

41 Shore J., Holland B.R., Sumner J.G., Nieselt K., Popinga A., and Wills □ P.R. (2019) Substitution Matrices Recapitulate Amino Acid Specificity of AARS Phylogenies. Mol Biol Evol Submitted.

42 Wolfenden R., Lewis C.A., Yuan Y., and Carter C.W., Jr. (2015) Temperature dependence of amino acid hydrophobicities. Proc Nat Acad Sci USA 112 7484–7488.

43 Yakovchuk P., Protozanova E., and Frank-Kamenetskii M.D. (2006) Base-stacking and base-pairing contributions into thermal stability of the DNA double helix. Nucleic Acids Res 34, 564–574

44 Yang X.-L., Otero F.J., Ewalt K.L., Liu J., Swairjo M.A., Köhrer C., RajBhandary U.L., Skene R.J., McRee D.E., and Schimmel P. (2006) Two conformations of a crystalline human tRNA synthetase–tRNA complex: implications for protein synthesis. EMBO J 25, 2919–2929.

45 Yaremchuk A., Cusack S., and Tukalo M. (2000) Crystal structure of a eukaryote/archaeon-like protyl-tRNA synthetase and its complex with tRNAPro(CGG). Embo J 19, 4745–4758.

46 Zuker M. (2003) Mfold web server for nucleic acid folding and hybridization prediction. Nucleic Acids Research 31, 3406–3415.

